# Inference based decisions in a hidden state foraging task: differential contributions of prefrontal cortical areas

**DOI:** 10.1101/679142

**Authors:** Pietro Vertechi, Eran Lottem, Dario Sarra, Beatriz Godinho, Isaac Treves, Tiago Quendera, Matthijs Nicolai Oude Lohuis, Zachary F. Mainen

## Abstract

Essential features of the world are often hidden and must be inferred by constructing internal models based on indirect evidence. Here, to study the mechanisms of inference we established a foraging task that is naturalistic and easily learned, yet can distinguish inference from simpler strategies such as the direct integration of sensory data. We show that both mice and humans learn a strategy consistent with optimal inference of a hidden state. However, humans acquire this strategy more than an order of magnitude faster than mice. Using optogenetics in mice we show that orbitofrontal and anterior cingulate cortex inactivation impact task performance, but only orbitofrontal inactivation reverts mice from an inference-based to a stimulus-bound decision strategy. These results establish a cross-species paradigm for studying the problem of inference-based decision-making and begin to dissect the network of brain regions crucial for its performance.

## Introduction

In natural foraging behaviors, animals must continually choose between trying to exploit resources at their current location and leaving to explore another, potentially superior one, at the expense of a possibly costly travel period. Viewed from the perspective of optimal decision-making, the crucial question is when is it best to leave the current site for another one? According to the marginal value theorem, in order to maximize returns, an optimal forager ought to leave its current site when the immediate rate of reward drops below the average rate (Charnov, 1976). However, this elegant solution to the foraging problem only applies in deterministic environments (Kolling et al., 2014), in which both immediate and average reward rates are knowable to the agent. In a more realistic scenario—for example, where rewards are encountered probabilistically—the immediate reward rate is ill-defined and the marginal value theorem does not apply.

One widely-used and powerful approach to model decision making in dynamic, stochastic environments is reinforcement learning (RL) (Sutton and Barto, 2018). In RL, the values of different actions (such as leaving a foraging site or staying on) are continuously updated through trial and error, based on their outcomes, allowing agents to adaptively modify their preferences as conditions change. In its simplest form, model-free RL assigns each action with a value that is updated based on its immediate outcome, with no regard to the causal, and often hidden, structure that links actions to outcomes. While computationally efficient and consistent with a large body of experimental data on both Pavlovian and operant tasks (Eshel et al., 2016; Schultz et al., 1997), model-free RL is not the best available strategy in many situations. Consider, for instance, a lion that has just successfully captured prey. If the fact that in doing this it has most likely scared away all other animals is ignored, the lion may continue to hunt in the same region, wasting a considerable amount of time searching for the now long-gone prey. Conversely, things may turn out badly for a zebra if it assumes that its current foraging ground was safe (that is, lion-free) just because it had not seen a lion yet in its immediate surroundings. What these examples illustrate is that relying solely on recent outcomes, while ignoring causal structures in the world, may have suboptimal (if not catastrophic) consequences. Instead, in structure learning (Boyen et al., 1999; Braun et al., 2010; Pearl, 1991), a form of inference-based RL, agents choose actions based on their beliefs about the current state of the world, which is determined by both incoming sensory evidence (such as outcomes) and knowledge of the underlying causal structure of the environment. How humans and animals implement such strategies remains an important and poorly understood question (Daw et al., 2011; Niv et al., 2015; Starkweather et al., 2018).

In the brain, it has been suggested that the orbitofrontal cortex (OFC) is crucial for hidden state representation, and hence for inference-based decisions. For example, in both rats and primates, lesions or inhibition of OFC impairs subjects’ ability to adjust their behavior in reversal learning tasks, where the depletion of a previously rewarding site (or the futility of a previously rewarding action) may be viewed as a change in the (hidden) state of the world (Wilson et al., 2014). The adjacent anterior cingulate cortex (ACC) on the other hand has been implicated in monitoring value during foraging (Hayden et al., 2011; Kolling et al., 2012) and could be responsible for encoding the value of alternative options (Kolling et al., 2016) and changing behavior based on the decreasing value of the current option (Shima and Tanji, 1998; Williams et al., 2004).

Here, we describe a foraging task in which subjects may seek rewards at either one of two foraging sites. This task has a special hidden structure: at any given moment, only one of the sites can deliver rewards and the side of the rewards switches with a certain probability after each foraging attempt. Importantly, even when reward is available, it is not delivered for every attempt, but rather with a probability less than 1. This makes the task a partially observable Markov decision process (POMDP): the true state of world (i.e., the identity of the rewarding site) is hidden and subjects must infer it based on noisy observations. A defining feature of this task, due to the hidden structure, is the asymmetry of the evidence provided by rewards and failures (unrewarded attempts): a single failure provides partial evidence in favor of a site switch, whereas a single reward provides full certainty that the current site is rewarding. A “stimulus-bound” agent, in the sense of Wilson et al. (2014), would assign value to observable states (being on the left or on the right or the other foraging site) by linearly combining rewards and failures. Such a process does not capture the essential asymmetry of the task. Ten rewards are much better than one reward in terms of value, but under optimal inference, one single reward is as informative as ten, since it already gives absolute certainty that the current site is active. Thus, leaving decisions under stimulus-bound and inference-based strategies in this task will be qualitatively different. A stimulus-bound agent will become more persistent the more rewards it has received at a site, whereas an inference-based agent will not show such an effect. We found that both mice and humans display hallmarks of inference in the performance of a foraging task and are able to build a non-trivial representation of task space. We further show that optogenetic inhibition of the OFC in mice selectively disrupts optimal inference behavior, biasing mice towards a sub-optimal stimulus-bound strategy. Similar inhibition of the adjacent anterior cingulate cortex (ACC), results in delayed leaving decisions but does not disrupt the inference process itself, suggesting a specific role of OFC in this important cognitive function.

## Results

### A probabilistic foraging task can dissociate value or evidence accumulation

We trained mice to perform a probabilistic foraging task (Fig. 1a). Subjects sought water rewards (2 µl each) by actively probing a foraging site. Each try at the active site yielded reward with probability *p*_*RWD*_, and could cause a switch with probability *p*_*SW*_(Fig. 1b). After a state switch, to obtain more rewards, subjects needed to travel to a second site at some distance and therefore bear a travel cost. Subjects were thus tasked with inferring a hidden state of the current site through a sequence of observations of stochastic events (rewards and failures). There are at least two distinct ways of integrating rewards and failures to form a decision. In a stimulus-bound process, the relative value of the left location with respect to the right location *v* = *v* _*LEFT*_ *-v*_*RIGHT*_ would increase gradually with left rewards, decrease gradually with right rewards, and decay to 0 with failures (Fig 1c.). In formulas, given a decay coefficient*γ*, a reward indicator *r*_*t*_, a side indicator *s*_*t*_ (1 for left, −1 for right) and signed outcomes: *o*_*t*_ *= r*_*t*_· *s*_*t*_

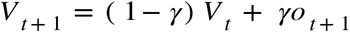

On the other hand, an agent that is aware of the structure of the task—the fact that a hidden state determines which side is rewarding at any time— could use rewards and failure to infer whether the current side is active or inactive. The relative value would then be:

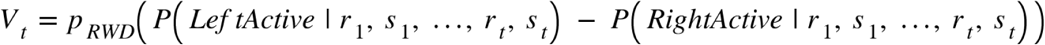

(see methods for a detailed treatment of the probability computation). Unlike the stimulus-bound mechanism in Fig. 1c, this process is able to track effectively the rapidly evolving value of the foraging sites (Fig. 1d). Both accumulation processes can be used as generative models of the behavior by defining the probability of staying on, e.g., the left side as a sigmoidal function of the relative value:

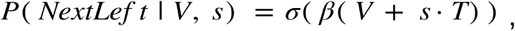

where *β* represents a softmax parameter (the higher it is, the more deterministic the behavior), and *T* represents the staying bias (when the value of the left and the right side are estimated as equal, the subject should still prefer to stay to avoid the travel cost). Both models predict that the probability of leaving increases with the number of consecutive failures. However, the effect of a reward is very different between them. In a stimulus-bound model the probability of leaving decreases with the number of rewards, as each reward contributes to the accumulated value. In the inference-based model it does not, as a single reward is sufficient to deduce with certainty that the current side is active (Fig. 1e). Thus a simple test of whether subjects are using inference is to check whether the number of failures before leaving changes with the number of preceding rewards (Fig. 1f).

**Fig. 1.**
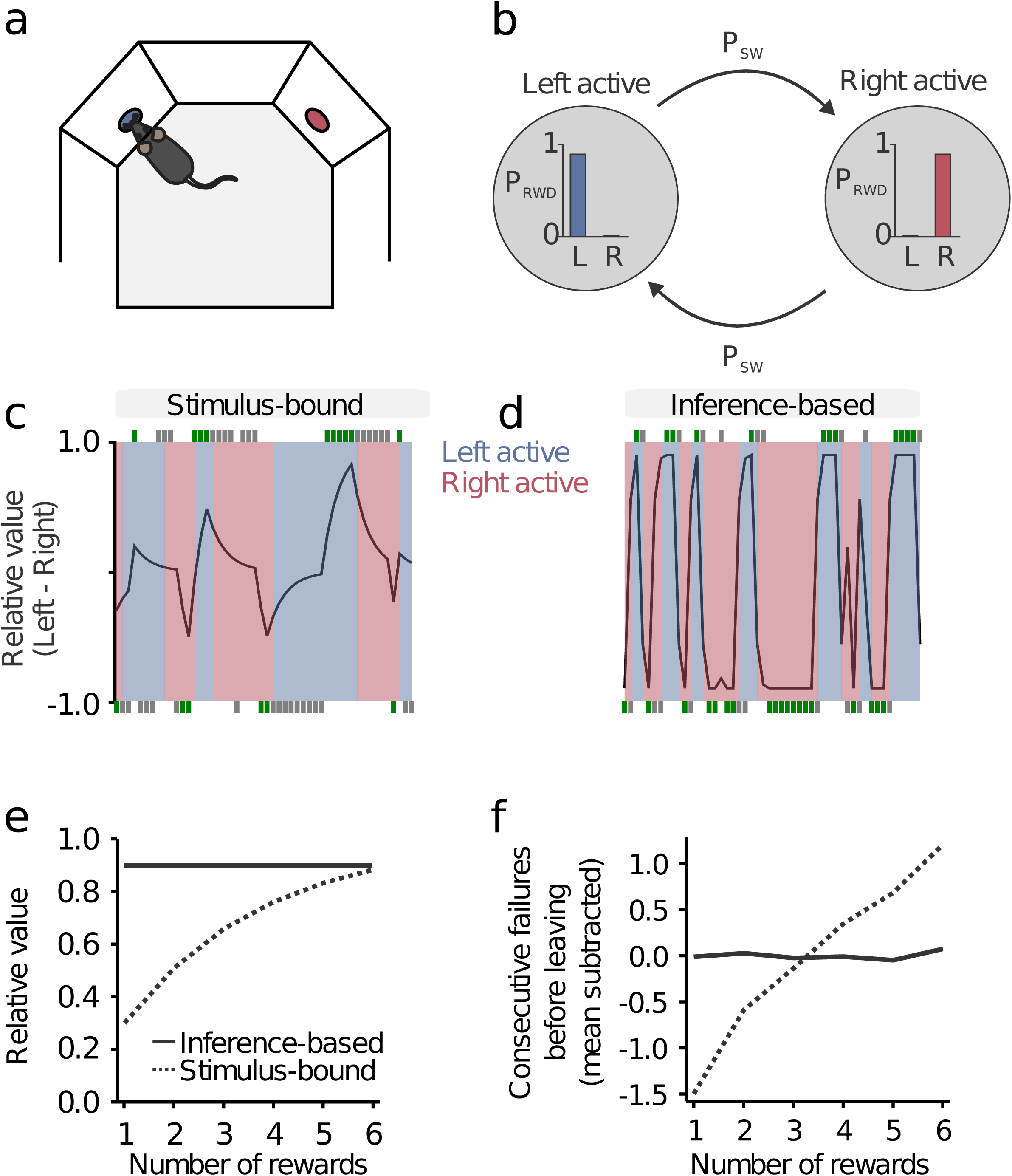
A probabilistic foraging task can dissociate stimulus-bound from inference-based evidence accumulation. **a** Schematic of rodent task. Mice shuttle back and forth between two reward sites to obtain water rewards. **b** Formally, the task is a hidden Markov model with *Left tActive* and *RightActive* states. It has two parameters: probability of reward given state and probability of state transition. **c, d** Estimated relative value (left minus right) as a function of trial history (rewards in green, failures in gray) in the stimulus-bound model and inference-based model respectively. Shaded patches indicate actual state. **e** Effect of rewards on relative value in stimulus-bound and inference-based models: the two models are simulated in a trial with only rewards on the same side. Relative value increases with reward number in the stimulus-bound but not in the inference-based model. **f** Consecutive failures before leaving (normalized subtractively) as a function of reward number in a simulated data of stimulus-bound and inference-based models: reward number has an effect on consecutive failures in the stimulus-bound but not in the inference-based model. See also Video S1.

### Mice accumulate evidence and not rewards

We trained 18 C57BL/6 wild type mice of 2 months age for 12 days in a baseline protocol with*p*_*RWD*_*=*0.9 and *p*_*SW*_= 0.3 and observed the effect of rewards on behavior during learning. In the early part of training, animals were unaware of the structure of the task and exhibited hallmarks of a stimulus-bound strategy: more failures were needed to leave the foraging port in rich foraging bouts, with many rewards before a state switch, compared to poor foraging bouts, with as little as one reward before a state switch. After training, however, the number of rewards had no effect on the number of failures before leaving, consistent with the inference-based model (Fig. 2a). To quantify this effect at a single animal level, we fitted a linear regression model that predicted the number of consecutive failures before leaving as a function of the number of prior rewards in the current foraging bout:*consecutiveFailures* ∼ 1 + *RewardNumber* (Fig 2b; here and throughout the text we use Wilkinson notation (Wilkinson and Rogers, 1973), see methods for a detailed explanation). The data show that during the first days of training there was a strong positive correlation between these two quantities, but with continued training this correlation decayed to zero (Fig. 2c). Therefore, experienced mice, unlike naive animals, decide when to leave the foraging site according to an inferred hidden state rather than directly integrating rewards and failures.

**Fig. 2.**
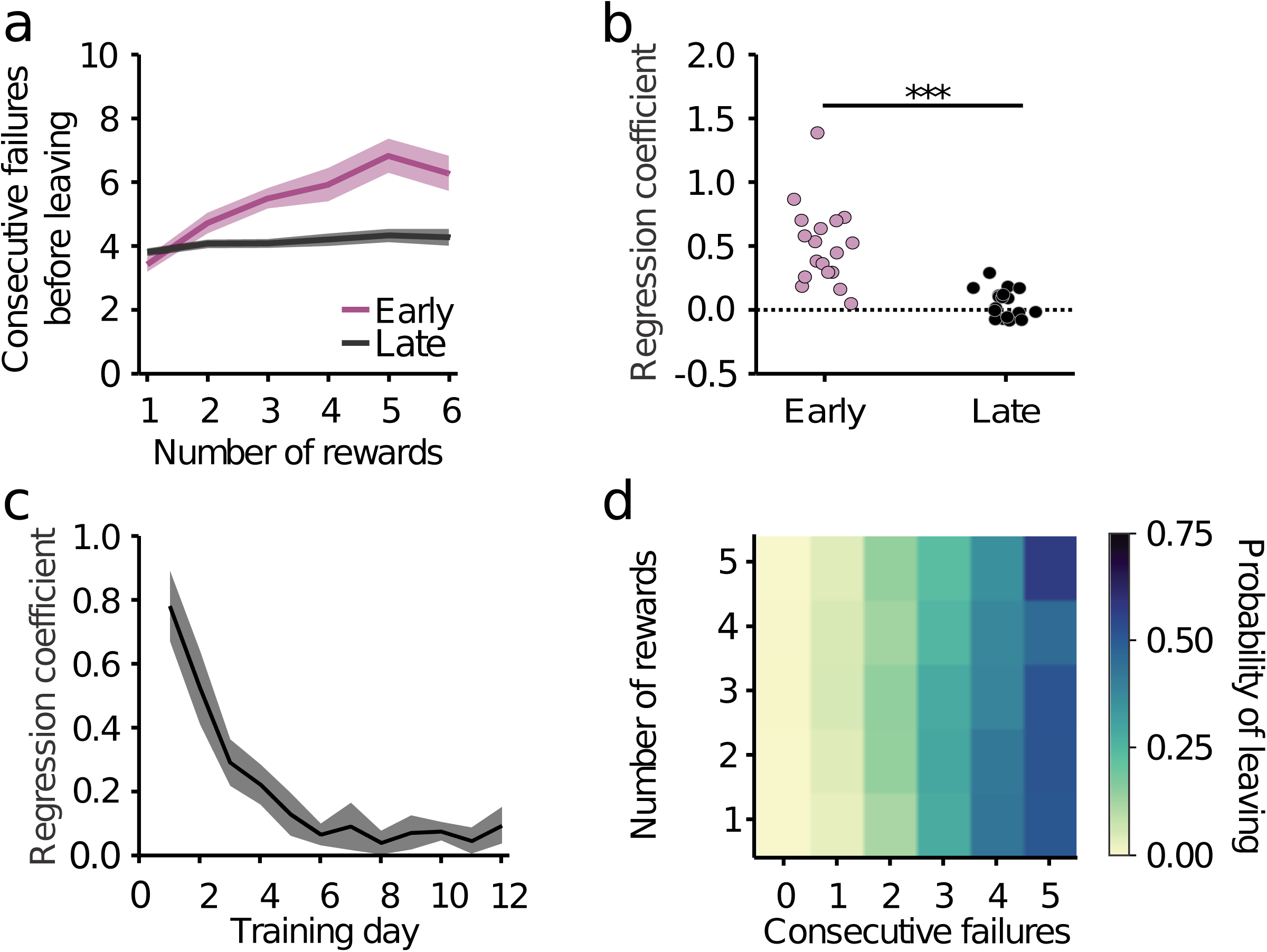
Mice accumulate inferred evidence for state switches and not port value. **a** Consecutive failures before leaving as a function of reward number in early training (day 1 to 3, purple) compared with late training (day 10 to 12, black). **b** Slope coefficient in *ConsecutiveFailures* ∼ 1 + *RewardNumber* for early training and late training. Slope coefficient is higher in early trials, likelihood ratio test on linear mixed-effect model *ConsecutiveFailures* ∼ 1 + *RewardNumber* + *Early* + *RewardNumber* + *Early* + (1| *MouseID*) versus a null model with no interaction: p < 1e-10, N = 18 mice (see methods for a description of the formula notation). **c** Evolution of reward number coefficient across days. **d** Probability of leaving as a function of number of rewards and consecutive failures in late training.

As another way of seeing this, a stimulus-bound integration strategy would weigh equally each reward and failure with opposite signs. Correct inference instead, given the structure of this task, requires that rewards are weighted nonlinearly (the first counting a lot and subsequent nothing) and differently from failures, which should add linearly. Indeed, in the trained mice, the effect of rewards and failures in shaping the behavior is qualitatively asymmetric in just this way, as can be seen by visualizing the probability of leaving as a function of both reward number and consecutive failures (Fig. 2d).

### Accumulation of evidence is tuned to task parameters

Having found that the foraging behavior of mice is consistent with the accumulation of evidence to infer a hidden world state, we asked whether this inference process is appropriately tuned to the statistics of the foraging environment, represented here by two parameters: reward probability *p*_*RWD*_ and state switch probability.*p*_*SW*_ Intuitively, if *p*_*RWD*_is high, then a single failure is strong evidence in favor of a state switch, leading to a faster accumulation process. Similarly, if *p*_*SW*_ is high, then a failure also carries more evidence in favor of a state switch compared to if it is low (Fig. 3a; see methods for a formal justification of this intuitive argument).

**Fig. 3.**
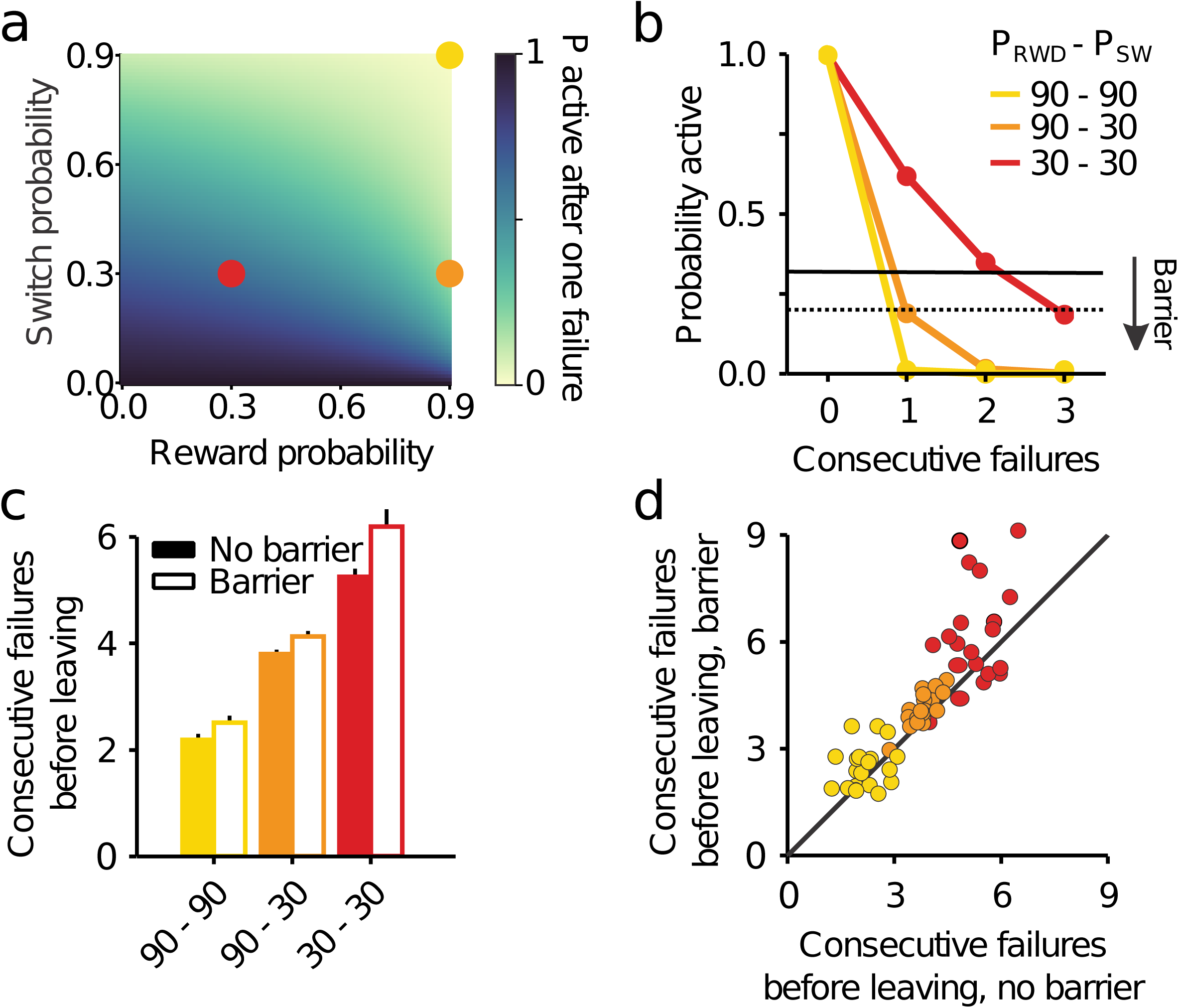
Accumulation of inferred evidence is tuned to task parameters. **a** Probability of being on the correct side after a failure as a function of reward probability and transition probability. **b** Probability of being on the correct side as a function of trial history for three protocols (Easy environment:*p*_*RWD*_ = 0.9 and *p*_*SW*_ = 0.9; Medium environment: *p*_*RWD*_ = 0.9 and *p*_*SW*_ = 0.3; Hard environment: *p*_*RWD*_ = 0.3 and *p*_*SW*_ = 0.3). Leaving decisions can be modelled by setting a threshold on this probability that changes as a function of the travel cost (black lines) **c** Consecutive failures before leaving as a function of the environment statistics and barrier condition. **d** Consecutive failures before leaving split by subject and environment statistics, barrier versus no barrier.

To test this, we trained a separate batch of mice on a set of three different foraging site statistics (Easy environment: *p*_*RWD*_ *= 0*.*9* and *p*_*SW*_ *= 0*.*3*; Medium environment: *p*_*RWD*_*= 0*.*9* and *p*_*SW*_ *= 0*.*3*; Hard environment: *p*_*RWD*_*= 0*.*9* and *p*_*SW*_*= 0*.*9*; and; see Fig. 3b). Since changing the foraging environment’s statistics can affect average reward rates (i.e. average number of rewards per trial), we adjusted the magnitude of individual rewards in order to equalize the amount of reward at a given site before state switch across conditions. As predicted normatively, mice increased the number of failed attempts they would tolerate as the state switching probability and the reward probability dropped (Fig. 3c, d; difference in failed attempts after last reward in Easy-Medium = −1.61 ± 0.03, difference Hard-Medium = 1.77 ± 0.04, N = 20 mice, likelihood ratio test on *consecutiveFailures* ∼ 1 + *protocol* + (1 | *MouseID*) versus *consecutiveFailures* ∼ 1 + *protocol* + (1 | *MouseID*) : p < 1e-10).

An important additional prediction of optimal decision theory in the context of a foraging task is that travel cost should modulate the threshold to leave a given foraging site. To test this, we increased the travel cost by placing a physical barrier between the two locations (travel time without barrier = 1.86 ± 0.13 s, N = 20 mice; travel time with barrier = 2.69 ± 0.13 s). Once again, the accumulation process was modulated consistently with the normative prediction, longer travel times resulting in a longer accumulation process and delayed leaving (Fig. 3c, d, effect of barrier in number of failed attempts after last reward = 0.42 ± 0.03, N = 20 mice, likelihood ratio test on *consecutiveFailures* ∼ 1 + *protocol* + *Barrier* + (1 | *MouseID*) versus *consecutiveFailures* ∼ 1 + *protocol* + (1 | *MouseID*) :p < 1e-10).

### Humans perform inference and tune behavior to task parameters

To test whether our findings were valid across species, we developed a translation of our behavioral assay for human subjects, in the form of a video game, where players would drag a character from one side of another of a touch screen and tap to achieve points (Fig 4a). The statistics of the video game (*p*_*RW*_ and *p*_*SW*_) were the same as those used in the rodent task.

**Fig. 4.**
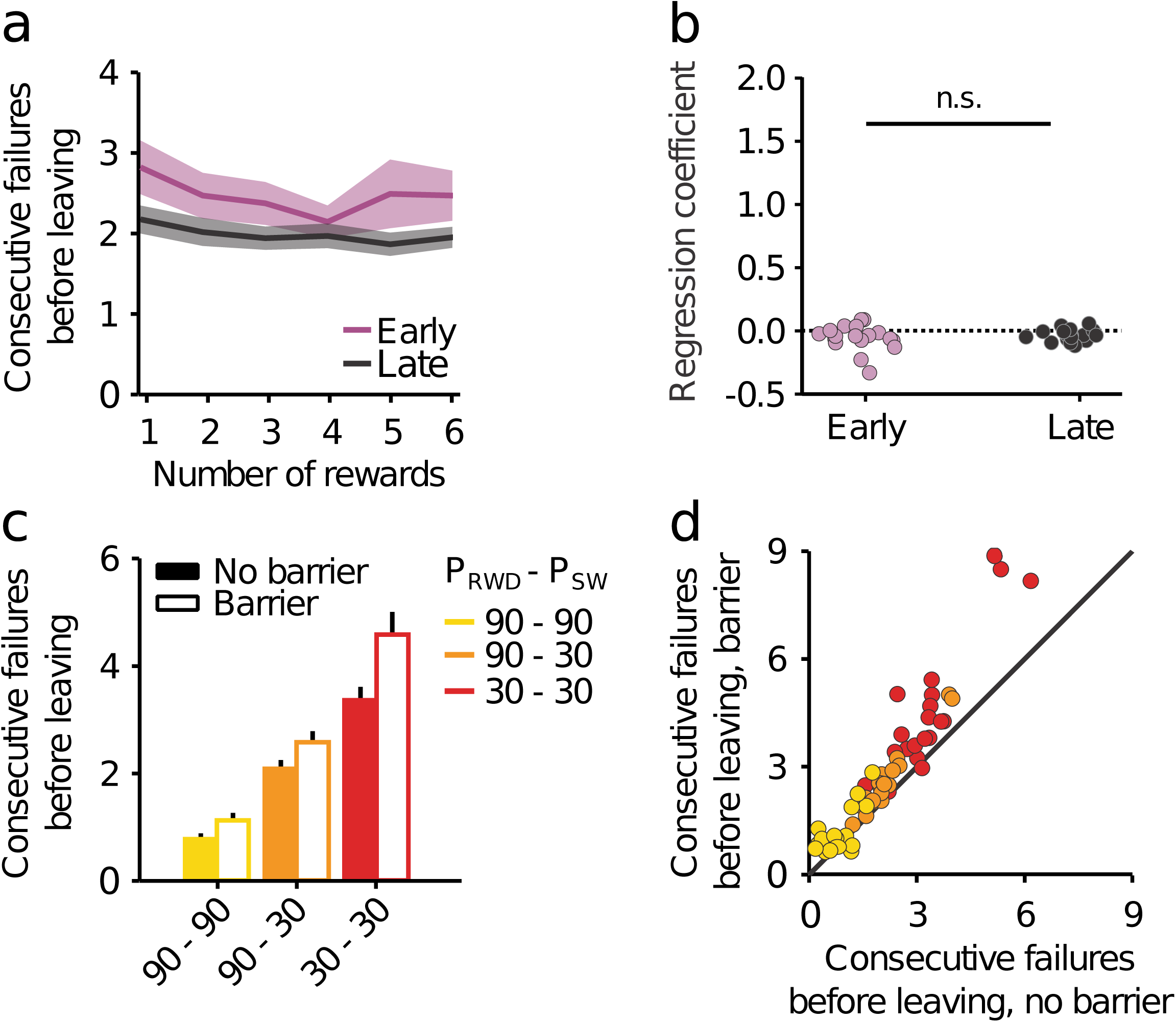
Humans perform optimal inference and tune behavior to task parameters. **a** The number of rewards has little effect on the probability of leaving during both early (purple) and late (black) training. **b** Number of consecutive failures as a function of reward number for human in early versus late part of training. Unlike mice, humans learn the statistics of the environment extremely quickly: slope coefficient is similar (and around 0) in both early and late trials: likelihood ratio test on linear mixed-effect model *ConsecutiveFailures* ∼ 1 + *RewardNumber* + *Early* + *RewardNumber* & *Early* + (1| *SubjectID*) versus a null model with no interaction: p = 0.45, N = 20 subjects. **c** Consecutive failures before leaving as a function of the environment statistics and barrier condition. **d** Consecutive failures before leaving split by subject and environment statistics, barrier versus no barrier. See also Video S2.

We again observed hallmarks of inference accumulation: the number of rewards had little to no effect on the behavior (Fig 4b, c), which is analogous to the behavior of the trained mice: unlike mice, however, humans needed almost no training to learn the statistics of the task and displayed hallmarks of evidence, rather than value, accumulation already in the first session.

Analogously to their rodent counterparts, human subjects modulated their behavior according to reward statistics as well as travel time (here affected by a manipulation in the character’s velocity) consistent with the normative predictions (Fig. 4d, e, difference in failed attempts after last reward in Easy-Medium = −1.39 ± 0.03, difference Hard-Medium = 1.48 ± 0.03, N = 20 subjects, likelihood ratio test on versus *consecutiveFailures* ∼ 1 + *protocol* + (1 | *subjectID*) versus *consecutiveFailures* ∼ 1 + (1 | *subjectID*) :p < 1e-10, effect of barrier in number of failed attempts after last reward = 0.59 ± 0.02, N = 20 subjects, likelihood ratio test on *consecutiveFailures* ∼ 1 + *protocol* +Barrier + (1 | *subjectID*) versus *consecutiveFailures* ∼ 1 + *protocol* + (1 | *subjectID*): p<1e-10).

### OFC, but not ACC, is necessary for the correct inference process

Finally, to study the brain mechanism of inference in this task, we tested the involvement of different regions of prefrontal cortex by silencing them using optogenetic stimulation of inhibitory GABAergic interneurons using VGAT-ChR2 mice (mice expressing channelrhodopsin-2 in GABAergic neurons). We examined 19 mice, of which 9 were bilaterally implanted with optic fibers (Supplementary Table 1) in the anterior cingulate cortex (ACC; Fig. 5a, S1a), 6 ChR2 expressing (HET) and 3 control wild type littermates (WT) not expressing ChR2 but implanted and stimulated in the same manner, and 10 were bilaterally implanted in the orbitofrontal cortex (OFC; Fig. 5a, S1b), 6 HET and 4 WT. Transient inactivation (3mW power per side, 10ms pulses at 75Hz, during poking in 50% of trials; Fig. 5b) of ACC significantly increased the average number of consecutive failures before leaving (Fig. 5c effect of stimulation on consecutive failures after last reward = 0.48 ± 0.05, N = 6 mice, likelihood ratio test on *consecutiveFailures* ∼ 1 + *protocol* + (1 | *MouseID*) versus *consecutiveFailures* ∼ 1 + *protocol* + (1 | *MouseID*) :p < 1e-10), while the same protocol applied to the controls had no effect (Fig. 5e, effect of stimulation = 0.003 ± 0.07, N = 7 mice, likelihood ratio test: p = 0.96). More specifically, we found that ACC inactivation multiplicatively increased the number of consecutive failures before leaving, consistently across protocols and animals (Fig. 5f,*protocol* and *stimulation* interact when predicting *consecutiveFailures*, likelihood ratio test: p = 1.46e-6, but not when predicting renormalized *consecutiveFailures*, likelihood ratio test: p = 0.15, N = 6 mice).

**Fig. 5.**
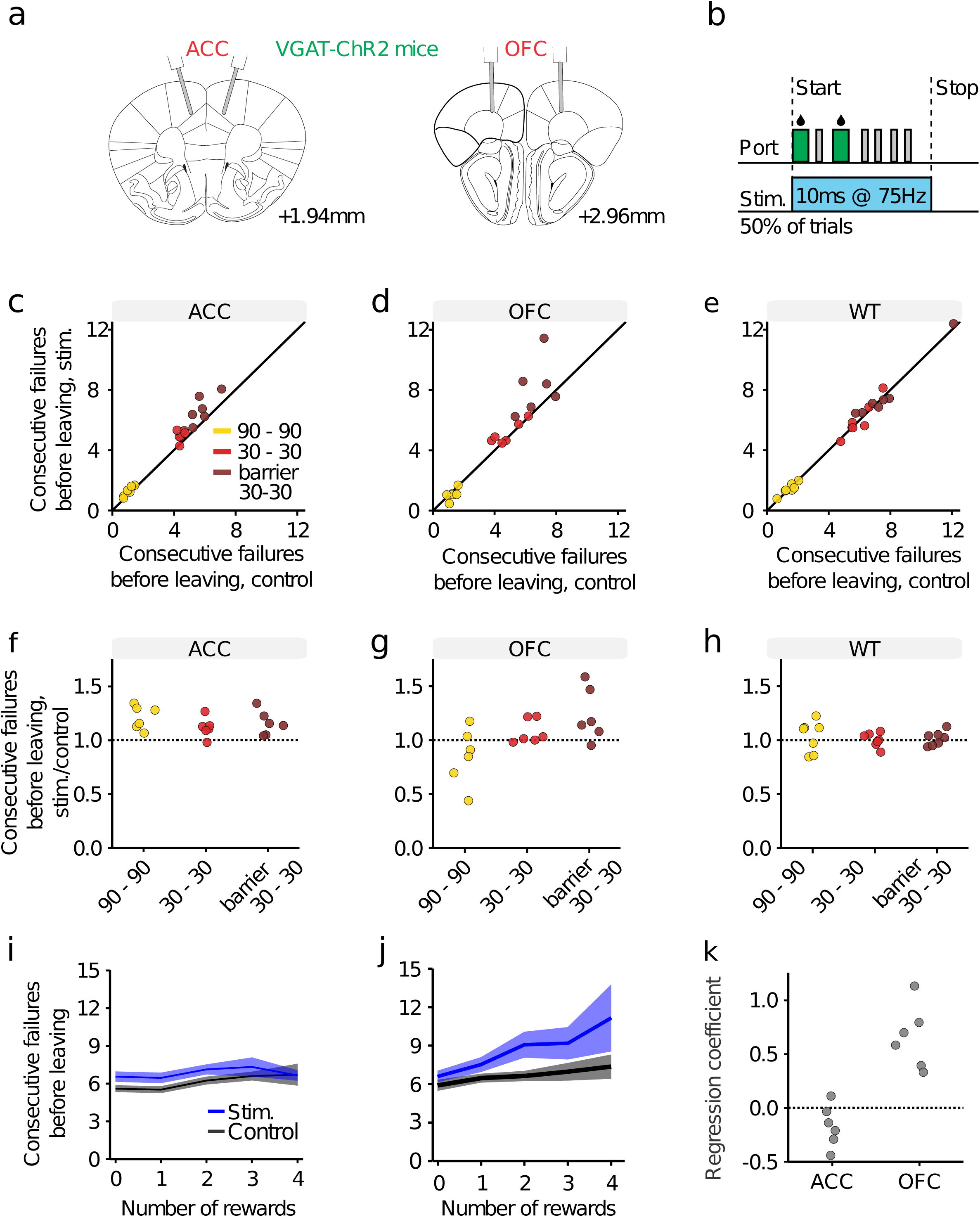
OFC, but not ACC, is necessary for optimal inference. **a** Scheme of the optic fiber placement. **b** Bilateral photostimulation at 3mW happened during nose-poking: it was triggered by the first poke in 50% of trials and lasted for 500 ms after the last poke in the trial. **c, d, e** Consecutive failures before leaving split by environment statistics, barrier condition and subject, inactivation versus control trials, for ACC implanted heterozygotes, OFC implanted heterozygotes and wild types, respectively. **f, g, h** Ratio of consecutive failures before leaving split in the same way as **c, d, e** for ACC implanted heterozygotes, OFC implanted heterozygotes and wild types respectively. When predicting renormalized consecutive failures,*stim* and *protocol* interact for OFC implanted heterozygotes (p < 1e-10, N = 6 mice) but not for ACC implanted heterozygotes (p = 0.15, N = 6 mice) or wild types (p = 0.77, N = 7 mice). **i, j** Number of consecutive failures as a function of reward number for ACC and OFC inactivation respectively (in the 30-30 protocol with barrier, which had the largest effect of ACC and OFC inactivation) **k** An animal by animal quantification: the coefficient of the interaction term in: *ConsecutiveFailures* ∼ 1 + *stimulation* + *reward number* & *stimulation* for ACC, distributed around 0, and OFC, greater than 0, in the 30-30 protocol with barrier. See also Figure S1 and Table S1.

Transient inactivation of OFC also increased the average number of consecutive failures before leaving (Fig. 5d; effect of stimulation on consecutive failures after last reward = 0.41 ± 0.08, N = 6 mice, likelihood ratio test on *consecutiveFailures* ∼ 1 + *protocol* + (1 | *MouseID*)versus *consecutiveFailures* ∼ 1 + *protocol* + (1 | *MouseID*) :p = 5.38e-7). However, unlike the case for ACC inactivation, it did so in an environment-statistics dependent manner. That is, the direction of effect for OFC inactivation actually reversed between easy and difficult protocols (effect of stimulation in hard protocol = 1.52 ± 0.24, effect of stimulation in easy protocol = −0.19 ± 0.05, N = 6 mice).

To investigate the involvement of these prefrontal areas in the inference process and task space representation, we considered a key difference in prediction between the inference model and the simpler stimulus-bound model, the effect of rewards on behavior. As noted above, under normal conditions, rewards fully reset the accumulation process, so that leaving times are not affected by the number of previous rewards (Fig. 1f). Strikingly, we found that OFC, but not ACC inactivation, disrupted this pattern: in OFC-inactivated trials, animals became sensitive to the number of rewards: the more rewards gained, the more delayed leaving decisions became (Fig. 5i, j, k, for ACC interaction term of stimulation and number of reward = −0.11 ± 0.13, N = 6 mice, likelihood ratio test: p = 0.37, for OFC, interaction term of stimulation and number of reward = 0.69 ± 0.22, N = 6 mice, likelihood ratio test: p = 0.002). This pattern of behavior is similar to the one observed in naïve mice first introduced to this task (Fig. 2a) and is indicative of an ineffective stimulus-bound strategy. Thus, the OFC has a special role in accurate state representation in a typical foraging environment in which states are hidden and require inference based on noisy observations.

## Discussion

In this study we developed a task in which subjects had to alternate between two foraging sites, only one of which was active at any given moment. The task embodied an important form of non-sensory uncertainty because the active port only delivered rewards with a certain probability. The task thus required subjects to infer whether each omitted reward was simply a stochastic failure or was instead an actual switch of state. To solve this task optimally, subjects were therefore required to infer a hidden state of the world (i.e. which site is active) rather than directly assigning a value to each foraging site. We found that, both mice and humans displayed hallmarks of optimal, inference-based behavior, reaching very similar solutions.

Our analysis of the behavioral data, particularly the number of consecutive non-rewarded tries before leaving, revealed that leaving decisions agreed with normative predictions of an inference-based foraging strategy in four important ways: (1) the number of consecutive failures was positively correlated with the propensity to leave; (2) rewards had a resetting effect on the leaving decision process; (3) subjects were sensitive to quantitative changes in the statistics of the foraging site; and (4) subjects were sensitive to the travel cost. However, mice and humans differed in an important way: while it took around six days for rodents to understand the environment statistics and integrate trial history correctly, humans started displaying hallmarks of the optimal behavior already during the first session. This difference may be due to faster learning but could also reflect the ability to generalize prior structural knowledge relevant to the task.

The accumulation of evidence is a primary cognitive computation. Similarly to sensory-guided tasks, in which integration of sensory evidence over time is needed to “average out” stimulus noise (Brunton et al., 2013; Gold and Shadlen, 2007; Shadlen and Newsome, 2001), here too repeated sampling is needed to determine which of two sites is currently rewarding. Specifically, each failure conveys ambiguous information, as it may be due to either an unlucky attempt at the rewarding site, or a guaranteed failure in the non-active site, and it is only by counting (integrating) the number of consecutive failures that a more accurate state estimation can be made. Our analysis of the leaving probability revealed that, much like in the sensory-based case, subjects do integrate this information when deciding whether to stay or leave. Moreover, by changing reward and transition probabilities, we were able to precisely control the amount of information associated with each failure, and observed that subjects readily adapted their leaving decisions to these changing conditions, such that the lower the information content of each sample was, the more such samples were needed before leaving.

In this task, the primary distinction between sub-optimal, stimulus-bound and optimal, inference-based strategies lies in the impact consecutive rewards have on leaving decisions. In a stimulus-bound behavior, which assigns values directly to the foraging site, the more consecutive rewards are gained at a given site, the higher the value of staying becomes, and consequently, leaving decisions tend to be delayed. In contrast, optimal inference in this task requires ignoring the number of consecutive rewards, since the delivery of a single reward is sufficient to know for certain which site is currently rewarding. As shown in Fig. 2, we found that, although initial behavior appeared to be sensory bound, after learning, subjects’ leaving decisions became independent of the number of rewards, consistent with an inference-based approach to leaving decisions.

Recent accounts (Schuck et al., 2016; Stalnaker et al., 2015; Wilson et al., 2014) proposed that the OFC is crucial for accurate state representations, particularly when states are hidden (that is, not explicitly given by the presence of a sensory cue, for example) and have to be inferred from fuzzy evidence. Our findings mesh well with this theory, as we found that OFC, but not ACC inhibition disrupted inference-based behavior. Unlike under control conditions, in which the number of failures before leaving was independent of the number of previously gained rewards, inhibiting the OFC resulted in mice performing more failures when experiencing large amounts of reward. This latter pattern is consistent with a stimulus-bound strategy, and therefore raises the intriguing possibility that the stimulus-bound strategy serves as a default behavioral approach, and is suppressed by the OFC when inference-based behavior is required.

The ACC has been implicated, in Pavlovian and operant tasks, as a potential candidate for the implementation of integration-to-threshold models. In Kawai et al. (2015), the authors observe neurons in the primate ACC whose firing rate scales with the number of consecutive negative outcomes in a Pavlovian task. From the foraging perspective, Hayden et al. (2011) reported cells in the ACC encoding the value of a depleting option. However, ACC inactivation in our task, unlike OFC inactivation, had only a modulatory effect on behavior: we did not observe qualitative changes in the strategy of the animals, but only an overall tendency to stay longer at the current port, which interacted multiplicatively both with the task statistics and with increased travel times. These results suggest that the ACC may encode the value of an alternative option, as proposed in Kolling et al. (2016), while not having a primary role in the inference process *per se*. By developing a human video-game and a rodent task requiring the same underlying computation to be solved, we could compare computational and cognitive processes across species. From a theoretical standpoint, this strengthens the generality of those results that held true for the two species, such as the ability to infer the hidden structure of the environment and to tune behavior to the environment-statistics. From a practical standpoint, the hidden state foraging task makes it possible to use rodent experimentation to guide human clinical research: the effect of drugs on cognitive quantities such as inference or persistence can be first tested in mice to then be validated in a clinical study. The more precise methods available in the rodent model could furthermore help dissect the mechanisms by which a specific drug affects cognition.

## Supporting information

Supplementary video (murine)

Supplemental video (human)

## Aknowledgments

We would like to thank Shira Lottem for her help with the graphic art of the human task. This work was supported by the European Research Council (Advanced Investigator Grants 671251 to Z.F.M.) and Champalimaud Foundation (Z.F.M.).

## Author Contributions

P.V., E.L., D.S., and Z.F.M. designed the experiments. P.V., D.S., B.G., I.T., T.Q. and M.N.o.L conducted the experiments. P.V. and E.L. analyzed the data. P.V. wrote the original draft. E.L., D.S. and Z.F. review and edited.

## Declaration of interests

The Authors declare no competing interests.

## MATERIALS AND METHODS

### EXPERIMENTAL MODEL AND SUBJECT DETAILS

#### Mice

Fifty-seven adult male C57BL/6 mice were used in this study. For the inference-based vs stimulus-bound behavior experiment (Fig. 2) 18 C57BL/6NCrl wild type mice of 2 months age were used. For the protocols manipulation experiment (Fig. 3), 20 wild type animal from different genetic backgrounds (8 Dat-Cre; 5 Gad2-Cre; 5 Sert-Cre; 1 VGAT-ChR2; 1 F512-Cre) of 6-8 months age were used, in order to reduce animal usage. For inactivation of anterior cingulate or orbitofrontal cortices (Fig. 5), 12 VGAT-ChR2 and 7 wild-type littermates were used. Mice genotypes were determined based on PCR and further verified using histological inspection of YFP expression which led to the exclusion of a single ACC implanted animal from further analysis (see Fig. S1c, d, e). The C57BL/6NCrl line was obtained from the Charles river laboratories, breeders were ordered and bred in-house for a maximum of 4 generations or 2 years (strain code: 027). The Dat-Cre mouse line was obtained from the Jackson laboratory (stock number: 006660). The Gad2-Cre was obtained from the Jackson laboratory (stock number: 010802). The Sert-Cre mouse line 61 was obtained from the Mutant Mouse Regional Resource Centers (stock number: 017260-UCD). The VGAT-ChR2 mouse line 8 was obtained from the Jackson laboratory (stock number: 014548). The FI12-Cre mouse line was obtained from the Mutant Mouse Regional Resource Centers (stock number: 017262-UCD). All experimental procedures were approved and performed in accordance with the Champalimaud Centre for the Unknown Ethics Committee guidelines and by the Portuguese Veterinary General Board(Direcao-Geral de Veterinaria, approval 0421/000/000/2016). The mice were kept under a normal 12 h light/dark cycle, and training, as well as testing, occurred during the light period. Before testing or after surgeries, for the inactivation experiments, mice were single-housed. During training and testing the mice were water deprived, and water was available to them only during task performance. Food was freely accessible to the mice in their home cages. Extra water was provided if needed to ensure that mice maintain no less than 80% of their original weight. For the protocols manipulation experiment behavioral training lasted 12 sessions, once per day, followed by 2 days of rest at the end of which we commenced testing. During training mice were exposed to the 3 different protocols (Easy: *p*_*RWD*_*=* 0.9 and *p*_*SW*_ *=* 0.9; Medium: *p* _*RWD*_*=* 0.9 and *p*_*SW*_ *=* 0.3; Hard: *p*_*RWD*_ *=* 0.9 and *p*_*SW*_*=* 0.9) for 4 consecutive days (1 day of adaptation and 3 of testing) before transitioning to the next environment. During testing, mice performed 1 session per day, 6 or 7 days a week. In protocols manipulation experiments, the sequence of protocols was counterbalanced across 2 groups of 10 mice (Group A: Hard, Medium, Easy; GroupB: Easy, Medium, Hard). In our analyses we considered 50 poke bouts per session after the first 10 during testing days and excluded poke bouts with no rewards.

#### Human participants

20 right handed healthy adults of Portuguese nationality (10 female and 10 male; 22 to 31 years of age), with no history of psychiatric diagnosis or prescribed drugs in the last 6 months, participated in this study. All participants gave written informed consent, and the study was conducted in accordance with the guidelines of the local ethics committee. The task consisted of 2 sessions of 1 hour, performed in different days with 2 to 10 days in between sessions. Each session consisted of 4 blocks with different protocols, and 10 minutes break after the second block. The sequence of protocols consisted of a block (Medium environment) followed by a short break (2 minutes), then a second block (Easy or Hard environment) followed by a long break (10 minutes), then a third block (Medium environment) followed by a short break (2 minutes) and a final block (Hard or Easy environment). The sequence of environments during testing was counterbalanced across 2 groups as described in mice experiments. In our analyses we considered all tapping bouts after the first 10 and excluded bouts with no rewards.

### METHOD DETAILS

#### Mice behavioural apparatus

The behavioral apparatus for the task was adapted from the design developed by Zachary F. Mainen and Matt Recchia (Island motion corporation, Nesconset, NY), originally developed for rat behavior. The behavioral box (15 × 12 × 18 cm, model 003102.0001, Island motion corporation), contained 3 front walls (135 degree angle between the center and the side walls) with 2 nose-poke port attached to the left and right front walls. For the inference-based vs stimulus-bound behavior experiment (Fig. 2), we used an acrylic custom made reproduction of the box (15 × 16 × 20 cm). Each port was equipped with infrared emitter / sensor pairs to report the times of port entry and exit (model 007120.0002, Island motion corporation). A nose-poke was considered valid if the infrared beam was broken for at least 100 ms. Water valves (LHDA1233115H, The Lee Company, Westbrook, CT) were calibrated to deliver a drop of 6 µl water for rewarded pokes in Easy and Hard environments and 2 µl of water in Medium environment: the reward size was adjusted to keep the reward amount per correct trial constant. The average number of rewarded attempts per correct trial *p*_*RWD*_/*p*_*SW*_ is, that is to say 1 in the easy and hard protocol (reward magnitude = 6 µl, amount of water per correct trial = 6 µl) and 3 in the medium protocol (reward magnitude = 2 µl, amount of water per correct trial = 6 µl). In optogenetic experiments, all protocols had an average of one reward per trial, but reward size was kept at 4 µl to increase trial number. In optogenetic experiments, blue LEDs were placed in the box ceiling and in all the ports to deliver a masking light. All signals from sensors were processed by Arduino Mega 2560 microcontroller board (Arduino, Somerville, US) and output from the Arduino Mega 2560 microcontroller board was implemented to control water and light delivery. Arduino Mega 2560 microcontroller was connected to the sensors and controllers through a Arduino Mega 2560 adapter board developed by Champalimaud Foundation Scientific Hardware Platform. An example behavioral video is available in the supplemental information.

#### Human video game task

Human subjects played a video game on a touchscreen device, with analogous features to the rodent behavioral assay. In the game, subjects control a character - a “witch” - on its quest to find and defeat an enemy. The witch must walk along the wall of a castle, shooting either the left or the right edge of this wall in search for the enemy that hides, at any given moment, in one of these two edges. The game obeys the same statistics as the rodent task: hitting the enemy is analogous to a water reward, the current location of the enemy corresponds to the active site, and every shot at the active side hits the enemy with probability.*p*_*RWD*_ Moving between the two sides of the wall has an associated cost (travel cost) that can also be manipulated with the appearance of rougher terrain (analogous to the physical barrier) that diminishes the traveling speed. As in the mouse case, reward size was manipulated to keep the average reward per correct bout constant (3 points). The game ended either when subjects collected 280 points or when a time limit of 20 minutes was exceeded.

Different environments had minor changes in the background images between them - for the medium protocol, since it was experienced twice per session, two different backgrounds were used. After the player transitioned to a different side, the enemy was displayed to cue whether the transition had been correct or if instead the player had to return to the previous side.

The human task was made using custom software developed using the game engine Construct2 (Scirra Ltd., Studio 117, The Light Bulb 1 Filament Walk Wandsworth, London, UK). Graphics were made by Shira Lottem and Tiago Quendera using Inkscape: Open Source Scalable Vector Graphics Editor. Audio assets were made by Tiago Quendera using Audacity(R) except for the Wilhelm Scream (Wikimedia Commons).

An example video (not from an experimental subject) showing the different environments is provided in the supplemental material. The task, open-source code and all assets are available at https://github.com/quendera/human-foraging.

#### Optogenetic stimulation

In order to optically stimulate ChR2 expressing VGAT-expressing GABAergic interneurons we used blue light from a 473 nm laser (LRS-0473-PFF-00800-03, Laserglow Technologies, Toronto, CA or DHOM-M-473-200, UltraLasers, Inc., Newmarket, CA) that was controlled by an acousto-optical modulator (AOM; MTS110-A1-VIS or MTS110-A3-VIS, AA optoelectronic, Orsay, FR) to deliver light 10 ms pulses of light at 75 Hz, connected to Arduino Mega 2560 microcontroller board (Arduino, Somerville, US). Light exiting the AOM was focused into an optical fiber patch cord (200 µm, 0.22 NA, Doric lenses Inc, 357 rue Franquet, Quebec, Quebec, CA), connected to a second fiber patch cord through a rotary joint (FRJ 1×1, Doric lenses), which was then connected to the chronically implanted optic fiber cannula (MFC_200/230-0.48_3mm_ZF1.25(G)_FLT; Doric lenses Inc, 357 rue Franquet, Quebec, Quebec, CA). We estimated an average 15% loss of light power between the patch cord tip and the optic fiber cannula before surgery. In order to deliver light at 3 mW power, previously to each experiment day, the laser power at the tip of the patch cord was adjusted to 3.6 mW, to account for the estimated power loss. To test each protocol, we habituated animals to the new protocol during two days, then stimulated during six consecutive days. Stimulation was delivered on 50% of trials and started with the first valid nose-poke. Stimulation ended if the animal did not nose-poke for 500 ms but would restart in case of another valid nose-poke on the same side.

#### Surgical procedures

Animals were anesthetized with isoflurane (4% induction and 0.5 - 1% for maintenance) and placed in a motorized computer-controlled Stoelting stereotaxic instrument with mouse brain atlas integration and real-time visualization of the surgery probe in the atlas space (Neurostar, Sindelfingen, Germany;http://www.neurostar.de). Antibiotic (Enrofloxacin, 2.5-5 mg/Kg, S.C.) and pain killer (Buprenorphine,0.1 mg/Kg, S.C.) and local anaesthesia over the scalp (0.2ml, 2% Lidocaine, S.C.) were administered before incising the scalp. Target coordinates were 1.9 mm A.P., ±0.5 mm M.L., 1.75 mm D.V. for ACC and 2.9 mm A.P., ±1.25 mm M.L., 1.8 mm D.V. for OFC. Two craniotomies were performed above the targets coordinates for OFC implants. For ACC implants fiber were implant over the target with an angle of ± 16° on the ML axis to avoid damage to the superior sagittal sinus, and two craniotomies were performed at coordinates 1.9 mm A.P.,±1 mm M.L.. An optical fiber (200μmcore diameter, 0.48NA, 510 mm) housed inside a connectorized implant (M3, Doric lenses, Quebec, Canada) was lowered into the brain (0°angle for OFC and 0°angle for ACC), through the craniotomy as the viral injection,and positioned10μmabove the target. The implant was cemented to the skull using Super Bond C&B(Morita, Kyoto, Japan) and once dried covered with black dental cement acrylic (Pi-Ku-Plast HP 36,Bredent, Senden, Germany). The skin was stitched at the front and rear of the implant. Gentamicin(48760, Sigma-Aldrich, St. Louis, MO) was topically applied around the implant. Mice were monitored until recovery from the surgery and returned to their home cage where they were housed individually.Gentamicin (48760, Sigma-Aldrich, St. Louis, MO) was topically applied around the implant. Behavioral testing started at least 1 week after surgery to allow for recovery.

#### Histology and microscopy

For histological analysis mice were perfused transcardially with 4% paraformaldehyde (PFA) in phosphate buffer solution (PBS). After removing the brain they were left for 24 hours in 4% PFA solution in PBS, then transferred in 0.1% sodium azide solution in PBS. Brains were sliced in 50 um coronal sections on a vibratome (Leica VT 1000 S), collected in wells maintaining the anterior-posterior order, and finally mounted in microscope slides (Thermo scientific, superfrost plus), with mowiol.

Fluorescent images were acquired with an automated slide scanner (AxioScan Z1) equipped with a 10×, 0.45 NA PlanApochromat objective and a Hamamatsu OrcaFlash camera. Use of the appropriate filter combination allowed for DAPI and EYFP acquisition (Beam Splitter: 395, excitation: 330-375, emission: 430-470, and Beam splitter: 498, excitation: 453-485, emission: 507-546 respectively).

Optic cannula placement was determined using coronal sections of the prefrontal cortex through which the fiber tract was visible. We determined the position by locating the section with the broadest base of the cannula tract and comparing the DAPI staining with the Allen Mouse Brain Atlas (Lein et al., 2007, Fig. S1,Supplementary Table 1).

#### Behavioral analysis and statistics

All analysis was performed using custom code written in Julia (Bezanson et al., 2017). The code used to simulate value or inference models is available on GitHub, under the MIT license, at https://github.com/piever/ValueInferenceTools.jl. The statistical analysis was performed using mixed effect models (McLean et al., 1991), in particular the Julia implementation MixedModels.jl (Douglas Bates et al., 2019). We fitted models with a random intercept (depending on subject identity) and compared nested models using a likelihood ratio test: in particular we used a chi-squared test on the difference of the deviance of the two nested models, using as many degrees of freedoms as the difference between the number of degrees of freedom of the two nested models (Wilks, 1938). That is to say, given two models *m* and *n* where is a special case of *m*:

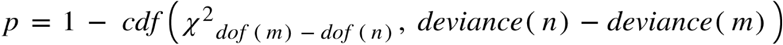

When the p value is too small, we do not report the value but simply write p < 1e-10, which is floating point notation for p < 10^-10^. To describe mixed models we will use Wilkinson notation (Wilkinson and Rogers, 1973), with | denoting random effects and & denoting interaction terms. For example the formula:

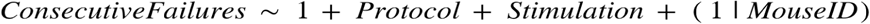

uses as predictor for the number of consecutive failures after last reward a constant intercept, a coefficient for each protocol different than the medium protocol (which we consider as baseline), a coefficient for stimulation and a random intercept across mice. The formula

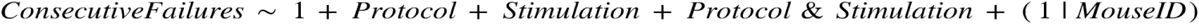

would also allow for an interaction term between protocol and stimulation.

#### Task design

We designed a probabilistic foraging task for parallel use in mice and humans. Subjects sought rewards (water or points, respectively), by actively probing a foraging site (nose-poking or screen-tapping, respectively). Each site could be in one of two states, active or inactive. Each try in the active state yielded reward with probability,*p*_*RWD*_ and could cause the site to switch to the inactive state with probability. *p*_*SW*_ This required the subjects to travel to a second, fresh, site at some distance and bear a travel cost. Subjects were therefore tasked with inferring a hidden state (active or inactive) through a stochastic sequence of observations (rewards and failures).

#### Relevant statistics in the task

After a rewarded attempt, the subject could be sure to be in the correct location: ambiguity comes from failures, as it was possible that the target was correct but the subject was being unlucky. The more unsuccessful attempts, the higher the probability of a transition having occurred. Accumulated evidence in favor of a switch is a monotonically increasing function of the task parameters *p*_*RWD*_ and *p*_*SW*_: the higher the reward probability, the more informative a failure is. Trivially, the higher the switch probability, the more likely the switch.

#### Possible task space representations

When analyzing our task, we consider two possible state representation. One is simpler and analogous to traditional approaches to modeling n-armed bandit tasks: the state corresponds to the current location of the subject (i.e. one of the two reward sites). The value of the two sites changes over time, yet the animals may be able to track this change with fast model-free learning. A second, more principled but more abstract approach, postulates that the subjects tries to infer the optimal state representation, i.e. the probability that their current location is rewarding. In this model, there is no longer any need for fast online learning as the task representation is stable. The computation happening in real time is the inference process to compute this probability. To account for variability in the behavior, we allow both decision noise, distributed according to the soft-max rule, as well as inference noise (the inference process may be suboptimal). We will refer to these two learning paradigm as Stimulus Bound Learning and Inference Based Learning respectively. It is important to note that, given the richer state representation, Inference Based Learning is the optimal way to solve the task and clearly outperforms simpler heuristics such as Stimulus Bound Learning.

#### Stimulus Bound Learning (SBL)

In Stimulus Bound Learning, we first define the relative valueas the difference of the value of the leftport and the value of the right port:

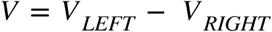

We defined two auxiliary variables: a reward variable *r* indicating the outcome of each reward attempts, i.e. 1 for a reward and 0 for a failure, and a side variable *s*, indicating the current side, 1 for left and −1 for right.

We can define a signed outcome *o* of each reward attempt, which is:

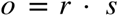

that is to say, 1 for a reward on the left, −1 for a reward on the right and 0 for an omission. For any attempt, we can then update the relative value using the signed outcome and some discount parameter *γ*

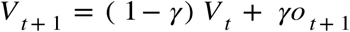

Which admits an explicit solution:

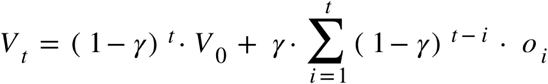

The probability of staying is a monotonically increasing function of the value of staying, so that rewards should make the animal more likely to stay and omissions more likely to leave, in a symmetric way.

#### Inference based learning (IBL)

We first derive recursive formulas to compute the probability that the current side is not rewarding as a function of the sequence of successful and failed reward attempts performed by the subject. In this model, the subject would compute the relative value as a function of the probability of the left (or right) side being active given the task history (*r*_1_,…,*r*_*t*_ represent the outcomes of the various attempts and *s*_1_,…,*s*_*t*_ the side of each attempt):

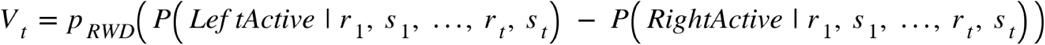

#### From value to decision

We have now defined to different ways to compute the relative value of left versus right, one directly based on reward accumulation, and one based on evidence accumulation. To define a behavior from this relative value, we need to consider two more parameters. First of all we need a bias term *T* : as the two foraging sites are far apart, subjects should prefer to repeat side rather than alternate, to avoid the travel cost. Then we need a “inverse temperature” parameter β to describe how deterministic the animal is (with a very high β the animal would almost always choose the option with greater value, whereas with β = 0 the animal would choose randomly). We can then use the soft-max rule to generate behavior:

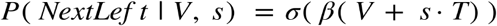

where *s* represents the current side (1 for left and −1 for right).

In the simulations we will use the same softmax rule to simulate behavior: difference between SBL and IBL derive from the different procedures used to compute the relative value.

#### Computing the likelihood ratio

In IBL we defined the relative value as a function of the relative difference:

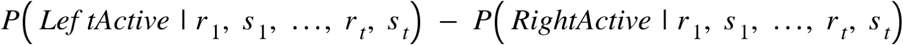

This quantity can be computed recursively. To do so we will need an auxiliary variable. We define *R*_*t*_ as the probability ratio that the current side is active or inactive given task history:

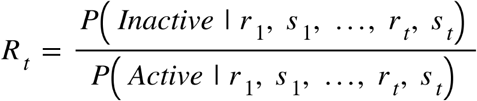

From *R*_*t*_ we can compute *V*_*t*_ as follows:

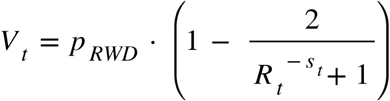

Rather than computing *V*_*t*_ recursively directly, we notice that *R*_*t*_ respects a simple recursive equation (likelihood ratio update equation):

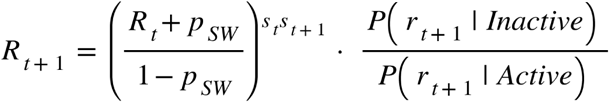

Where *p*_*SW*_ represents the probability of switching from active to inactive state.

The term 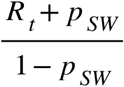 is the ratio between the following two equations:

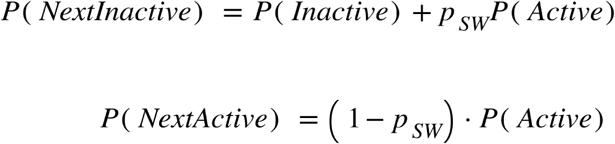

The exponent *S*_*t*_*S*_*t*+1_ simply means that the probability ratio 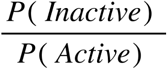 inverts when the subject changes side. Finally the term 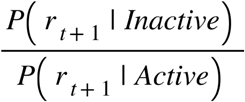 represents the new evidence acquired with the outcome of attempt *t* + 1.

In the case of a reward, *P*(*r*_*t*+1_ |*Inactive*) = 0, so:

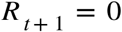

If *r*_t+1_is a failure, then *P*(*r*_*t*+1_ |*Inactive*) = 1 whereas *P*(*r*_*t*+1_ |*Active*) = 1 − *p*_*RWD*_, so the likelihood ratio update equation simplifies to:

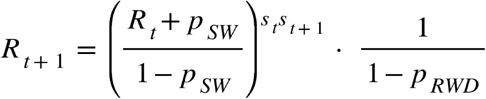

Having established that *R*_*t*_ resets to 0 with a reward, we can analyze the most interesting case for a probability computation: a sequence of attempts on the same side (let us say *s*_1_,…, *s*_*t*_ = 1) where the first attempt is rewarded (thus resetting the probability) and the following are not.

As the first attempt is rewarded, *R*_1_=0. Furthermore, the likelihood ratio, combined with the assumption that all attempts are on the same side, grows following the recursive equation:

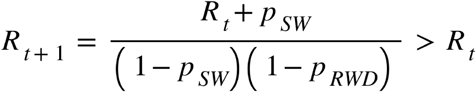

We can define the auxiliary quantity

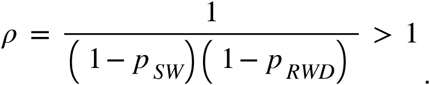

Our recursive equation becomes:

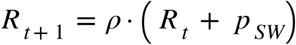

This is a standard linear recursion that we can solve with linear transformation

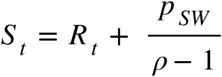

The recursion of *S*_*t*_ is:

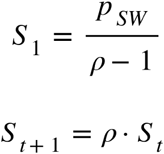

whose solution is

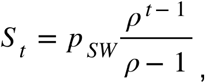

therefore:

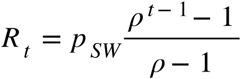

That is to say *R*_*t*_grows exponentially with rate log(*ρ*) = −log(1 − *p*_*RWD*_) − log(1 − *p*_*SW*_): increasing either *p*_*RWD*_ or *p*_*SW*_ would increase the growth rate of *R*.

**Supplementary Table S1.**
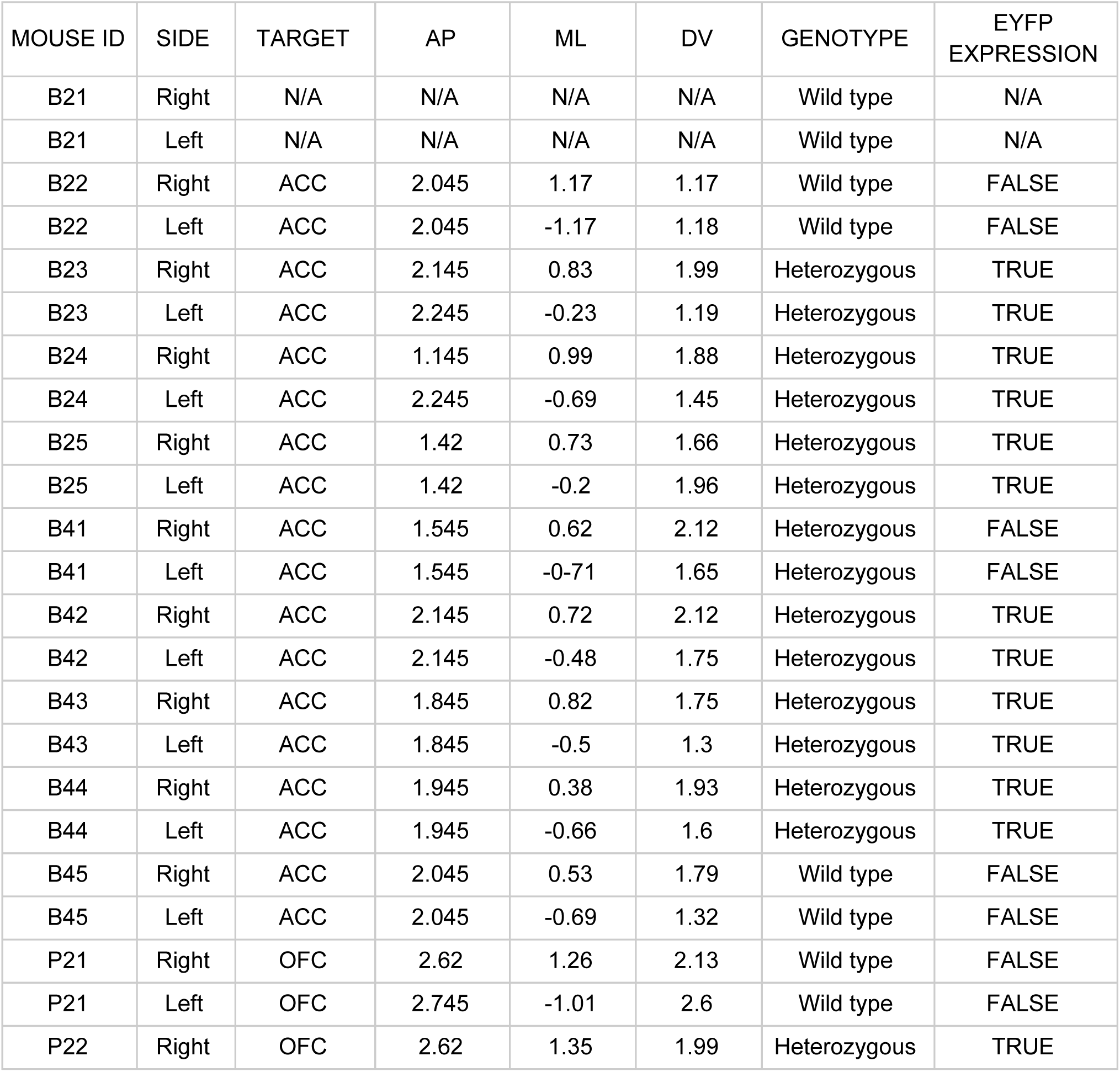
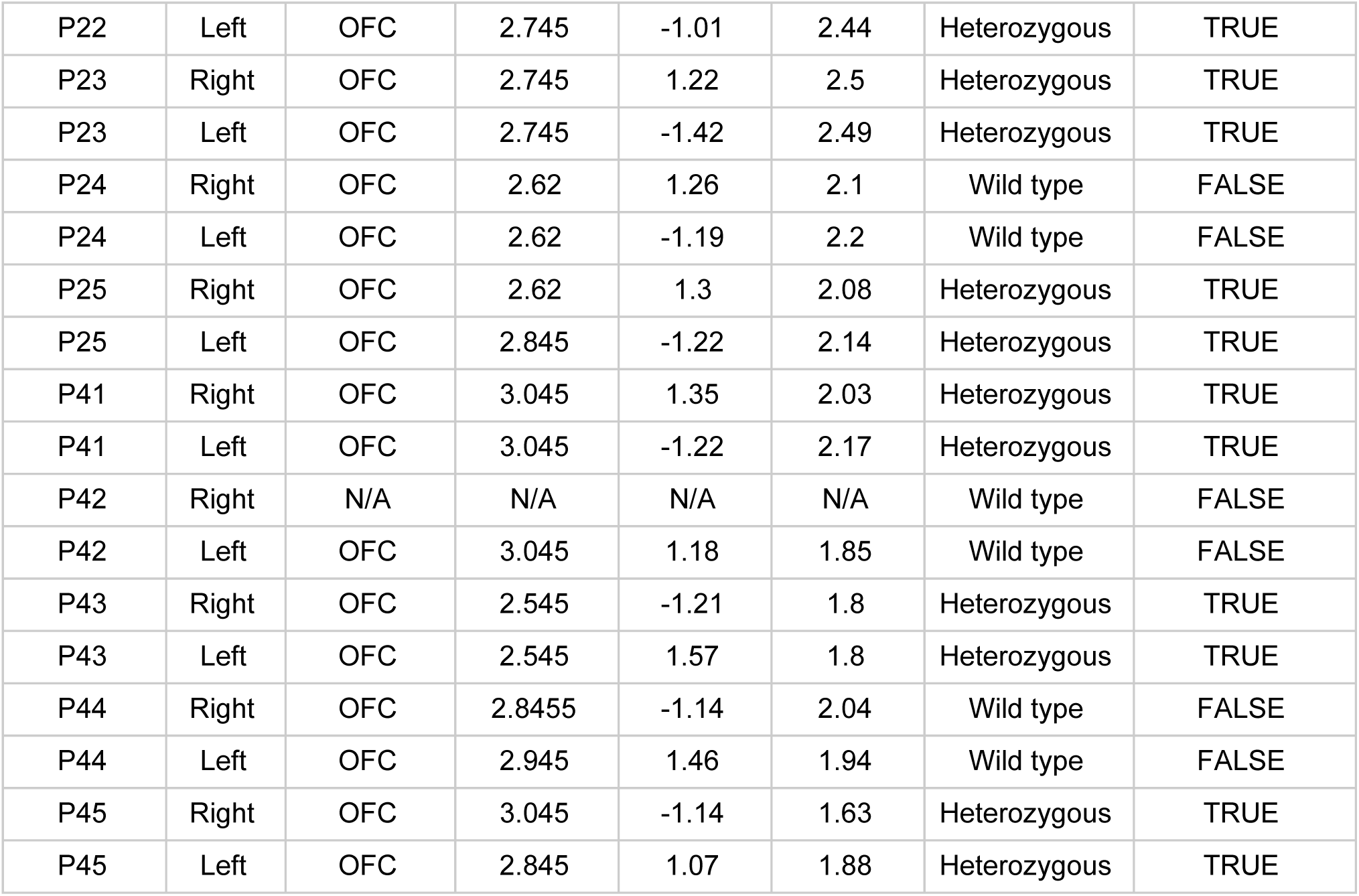
Related to Figure 5 and Supplementary Figure 1. Optic fibers placement coordinates and expression EYFP. Location of optic fibers across all VGAT-ChR2 animals used for this paper with anteriorposterior (AP), mediolateral (ML) and Dorsoventral (DV) coordinates. In mouse B21 wasn’t possible to perform histological controls, due to sudden death of the animal after the experiment period, that precluded from performing the perfusion of its brain. In mouse P42 wasn’t possible to determine the placement of the fiber on the right emisfere, due to damage in the slices during cutting. The expression of EYFP conjugated to ChR2, was asses through fluorescence widefield microscopy to confirm the animal genotype. In mouse B41 EYFP signal wasn’t detected (Sup.Fig1). despite the animal was initially genotyped as heterozygous and it was excluded from the analysis.

