## Supplemental information for "Inference based decisions in a hidden state foraging task: differential contributions of prefrontal cortical areas"

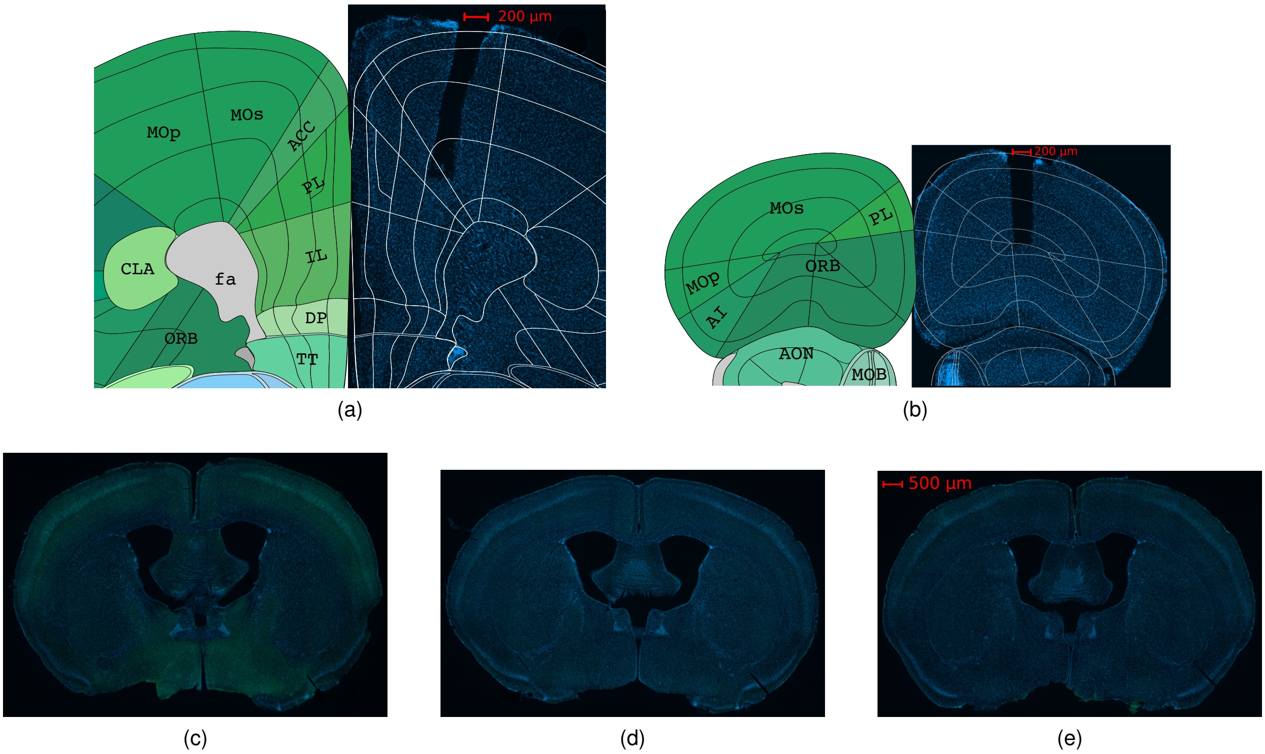


**Figure S1. Related to Figure 5.** Inhibition of ACC or OFC cortex in VGAT-ChR2 mice. **a, b** Coronal section from 2 VGAT-ChR2 mouse (blue, DAPI). Dark area show the location of the optic cannula over ACC in **a** or over OFC in **b**. Correct positioning of the fiber was verified overlapping reference images from the coronal Allen Mouse Brain Reference Atlas. ACC, anterior cingulate; AI, agranular insular area; AON, anterior olfactory nucleus; CLA, claustrum; DP, dorsal peduncular area; fa, corpus callosum; ILA, infralimbic area; MOp, primary motor area; MOs, secondary motor area; PL, prelimbic cortex; ORB, orbital area; TT, taenia tecta. Image credit: Allen Institute. **c, d, e** Fluorescence widefield microscopy of EYFP signal conjugated to ChR2 in mouse VGAT-ChR2-EYFP line 8, in an example heterozygote **c**, an example wildtype **d** and an animal initially genotyped as heterozygous that was excluded from the dataset due to lack of EYFP expression **e**.
